# Reference genome sequences of the oriental armyworm, *Mythimna separata* (Lepidoptera: Noctuidae)

**DOI:** 10.1101/2022.11.18.516534

**Authors:** Kakeru Yokoi, Seiichi Furukawa, Rui Zhou, Akiya Jouraku, Hidemasa Bono

## Abstract

Lepidopteran insects are an important group of animals, among which some are used as biochemical and physiological model species in the insect and silk industries, whereas others are major agricultural pests. Therefore, genome sequences of several lepidopteran insects have been reported thus far. The oriental armyworm, *Mythimna separata*, is an agricultural pest commonly used to study insect immune reactions and interactions with parasitoid wasps as hosts. To improve our understanding of these research topics, reference genome sequences were constructed in the present study. Using long-read and short-read sequence data, *de novo* assembly and polishing were performed, and haplotigs were purged. Subsequently, gene predictions and functional annotations were performed. To search for orthologs of the Toll and immune deficiency (IMD) pathways and C-type lectins, annotation data analysis, BLASTp, and Hummer scans were performed. The *M. separata* genome is 682 Mbp; its contig N50 was 2.7 Mbp with 21,970 genes and 24,452 coding sites predicted. All orthologs of the core components of the Toll and IMD pathways and 105 C-type lectins were identified. These results suggest that the genome data were of sufficient quality as reference genome data and could contribute to promoting *M. separata* and lepidopteran research at the molecular and genome levels.

**Simple Summary:** The oriental armyworm, *Mythimna separata*, an agricultural pest, is commonly used to study insect immune reactions and interactions with parasitoid wasps. To promote such studies, a reference genome was constructed. The *M. separata* genome is 682Mbp long—a size comparable to that of other lepidopteran insects. The contig N50 value of the genome is 2.7 Mb, which indicates sufficient quality to be used as reference genome data. Gene set data were constructed using genome and RNA-sequencing data; a total of 21,970 genes and 24,452 coding sites were predicted. Functional gene annotation was performed using the predicted amino acid sequences and reference gene set data of the model organism and other insect species as well as Unigene and Pfam datasets. Consequently, 45–80% of the amino acid sequences were annotated using these data sets. Using these data, most of the orthologs of core components in the Toll and immune deficiency (IMD) pathways were identified, suggesting the presence of these two pathways in *M. separata*. Additionally, 105 C-type lectins were identified in the *M. separata* genome, which were more numerous than those in other insect species, suggesting that these genes may be duplicated.

## 1. Introduction

Several species of lepidopteran insects are known, of which some are beneficial and some detrimental. Although the domestic silkworm (*Bombyx mori;* Bombycidae; Linnaeus, 1758), is used for silk production, some lepidopteran insects are major agricultural pests. For example, the fall armyworm (*Spodoptera frugiperda;* Noctuidae; J. E. Smith, 1797), the beet armyworm *(Spodoptera exigua*; Noctuidae; Hübner, 1808), the tobacco cutworm (*Spodoptera litura*; Noctuidae; Fabricius, 1775), The diamondback moth (*Plutella xylostella;* Plutellidae; Linnaeus, 1758), the cabbage looper (*Trichoplusia ni*; Noctuidae; Hübner, 1800–1803), and the oriental armyworm (*Mythimna separata;* Noctuidae; Walker, 1865) are the most severe pests in some crops [1–3]. In contrast, the tobacco hornworm (*Manduca sexta;* Sphingidae; Linnaeus, 1763) has been used as a biochemical and physiological model species of insects (e.g., to study immune reaction, development, and metamorphosis [4,5]). Because of this importance, research on the lepidopteran family of insects is quite active, and the genomes of some lepidoptera species have been sequenced. The first lepidopteran genome sequence data to be reported was the draft genome sequence data of *Bombyx mori* in 2004 [6,7]. The genome data was updated [8] and, the chromosome-level genome sequence of *B. mori* was reported [9]. Additionally, genome sequence data of other lepidopteran families, including *S. frugiperda* [10,11], *S. exigua* [12], *S. litura* [2], *P. xylostella* [13,14], *T. ni* [15], and *M. sexta*, have been reported [16,17]. In addition, the genome sequence data for several butterflies were reported [18].

*M. separata* is commonly used to study immune reactions in insects. Insects exhibit humoral and cellular immune responses to foreign microorganisms [19]. *M. separata* larval hemocytes, especially granular cells and plasmatocytes, play a central role in cellular immune reactions, and different types of invaders activate different reactions such as encapsulation, phagocytosis, and nodule formation. Several molecules in the *M. separata* hemolymph mediate these reactions. Growth blocking peptides, found as insect cytokines, trigger the spreading of plasmatocytes [20]. C-type lectins (CTLs) are a superfamily of proteins that recognize carbohydrates in a calcium-dependent manner. Some CTLs can activate cellular immune reactions. Ishihara et al. (2017) reported that the CTL encapsulation promoting lectin (EPL) enhances encapsulation [21]. Another CTL, IML-10, also promotes encapsulation [22]. A transcriptome analysis found 35 CTLs from *M. separata larvae* [23], and many of them changed their expression profiles upon bacterial exposure, suggesting that they are involved in immune regulation. To study the interactions between *M. separata* and parasitic wasps or flies, the oriental armyworm is used as a host insect for these species [24–28]. Using both species, our group identified several candidate factors needed for successful parasites in the braconid wasp, *Meteorus pulchricornis* (Hymenoptera: Braconidae; Haliday, 1835) [29] and investigated several gene functions of *M. separata* related to apoptosis or immune reactions considered to be essential for parasitoids [30,31].

To deepen our understanding of the molecular mechanisms of immune reactions and interactions between *M. separata* and parasitoid wasps, we constructed a reference-quality genome sequence of the oriental armyworm (*M. separata*), using long-read and short-read sequence data. Consequently, gene set data and the translated amino acid sequence data were prepared, and functional annotations of the predicted genes were performed (Figure 1). Using the gene set data, the orthologues consisting of IMD and Toll pathways, which are major immune signaling pathways, were searched. Finally, CTL genes were searched in the gene sets, and sequence and domain analysis of CTL were performed. These results suggested that the reference genome data and gene set data could contribute to promoting the *M. separata* and Lepidopteran research at the molecular or genome level.

**Figure 1.**
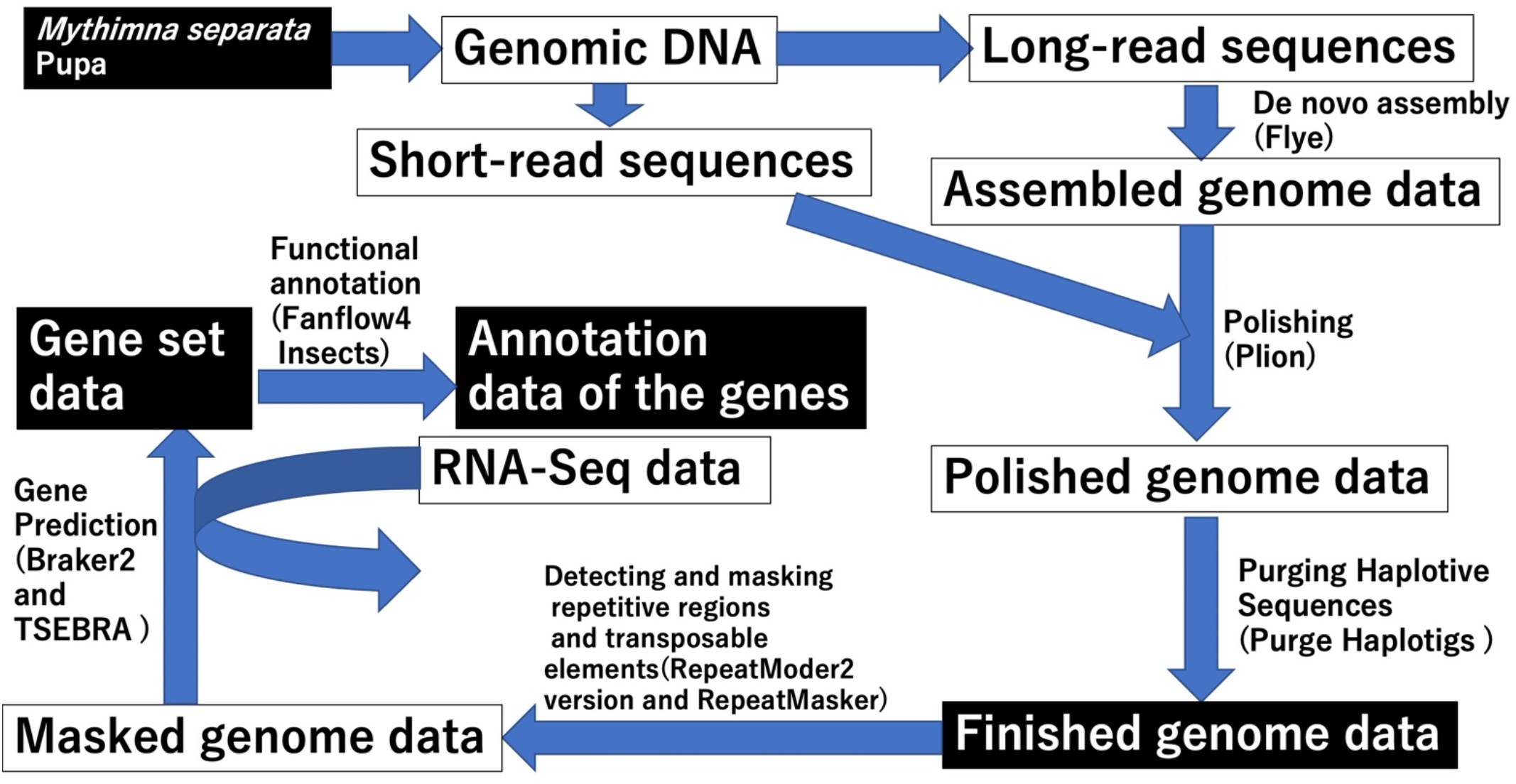
Schemes of the reference genome constructions, gene prediction and functional annotating. “Finished genome data” indicates the reference genome data. The names of software used in each step are shown in brackets

## 2. Materials and Methods

### 2.1. Sample preparation and sequencing

*M. separata* was supplied from stock cultures stored at Takeda Chemical Industries, Ltd. [32] and maintained in the Laboratory of Applied Entomology and Zoology, University of Tsukuba, Japan. The insect was reared on an artificial diet (Silkmate, Nihon Nosan Kogyo, Kanagawa, Japan) at 25±2 ^°^C, 40–80% relative humidity (r. h.), and an L16:D8 photoperiod.

Genomic DNA from single male pupae was extracted using NucleoBond HMW DNA (TaKaRa Bio Inc, Shiga, Japan) according to manufacturer’s protocol. Briefly, the whole body in 500 µL Lysis buffer was homogenized with a plastic pestle and liquid nitrogen. After treatments of 200 µL Liquid Proteinase K at 50 °C for 2 h and 100µL Liquid RNase A at 25 °C for 5 min, the homogenate was loaded onto NucleoBond HMW Column. Through washing step, DNA was finally eluted with 5 µL Elution buffer. Prep Kit 2.0(Pacific Bioscience, CA, USA) according to manufacturer’s protocol. The sequencing library was size-selected using the BluePippin system (Saga Science, MA, USA) with a lower cutoff of 30 kbp. One SMRT Cell 8M was sequenced on the PacBio Sequel II System with Binding Kit 2.0 and Sequencing Kit 2.0, yielding a total of 6,971,329 polymerase reads (201,369,397,241 bp). Shearing System M220 (Covaris Inc., MA. USA). A paired-end library was constructed with a TruSeq DNA PCR-Free Library Prep kit (Illumina, CA, USA) and was size-selected on an agarose gel using a Zymoclean Large Fragment DNA Recovery Kit (Zymo Research, CA. USA). The final library was sequenced on the Illumina NovaSeq 6000 sequencer with a read length of 150 bp.

Total RNA samples were prepared from the hemocytes of *M. separata* larvae, using TRIzol (Thermo Fisher Scientific, Waltham, MA, USA). PolyA RNA libraries were constructed using the TruSeq Stranded mRNA Library Prep Kit (Illumina), and sequencing (paired-end sequencing with 100-nt reads) was performed on a NovaSeq6000 platform (Illumina).

All raw sequence data were deposited in the Sequence Read Archive (SRA) of the DNA Data Bank of Japan (DDBJ). SRA accession IDs of the raw sequence data used in this study are listed in Supplementary Data 1.

### 2.2. Genome assembly and gene prediction

The output BAM file of Sequel II was converted to FASTA file using BAM2fastx version 1.3.1 (URL: https://github.com/PacificBiosciences/bam2fastx, accessed on 2 September 2022). *De novo* assembly was performed using the converted FASTA file with Flye version 2.9-b1774 with default settings [33]. Polishing of the assembled genome was performed using Minimap2 version 2.17 (mapping) [34], SAMtools version 1.10 (file conversion) [35], and Plion version 1.23 (perform polishing) [36]. Identification of the allelic contig pairings and productions haplotype-fused assemblies were performed using Purge Haplotigs version 1.1.2. The genome sequences was assessed using BUSCO version5.2.2 with eukaryota_odb10 file [37]. Status of the genome sequences were evaluated using assembly-stats version 1.0.1 (URL: https://github.com/sanger-pathogens/assembly-stats accessed on 2 September 2022). The finished genome sequences were deposited to DDBJ/ENA/GenBank (accession IDs: BSAF01000001BSAF01000569; Supplemental Data 2)

To search for repetitive sequences and transposable elements (TEs), RepeatModer2 version DEV and RepeatMasker version 4.1.2-p1 were used [38,39]. Gene prediction was performed using masked sequences and RNA-sequencing (RNA-Seq) data, including those from the public database as hint data (Supplementary Data 1). RNA-Seq data mappings to the reference genome were performed using HISAT2 version 2.2.1 [40] and mapped data were converted using SAMtools. Gene predictions were performed twice using Braker2 version 2.1.6 with RNA-Seq data-hint only mode and protein data-hint only mode independently[41]. The two output data from RNA-Seq data-hint only mode and protein data-hint only mode were merged by TSEBRA version 1.0.3 [42]. The amino acid sequences of the gene sets were functionally annotated by the functional annotation workflow Fanflow4Insects [43]. The functional annotation included the assignment of the top hit genes from comprehensively annotated organisms using the sequence similarity search program in global alignment (GGSEARCH) (https://fasta.bioch.virginia.edu/; accessed on 1 August 2022) and the protein domains using the protein domain database Pfam (http://pfam.xfam.org/; accessed on 1 August 2022) via HMMSCAN in the HMMER package (http://hmmer.org/; accessed on 1 August 2022). Accession IDs of raw read sequences of the RNA-Seq is listed in Supplemental Data 1. To remove the contaminating microbe sequences, the predicted amino acid sequences were annotated using BLAST nr database (Downloaded via the URL: https://ftp.ncbi.nlm.nih.gov/blast/db/ accessed at March 3, 2022, 444,154,253 protein sequences.) by BLAST search (version 2.12.0+).

### 2.3. Searching for and sequence analysis of Immune-related genes in M. separata

To search for genes related to IMD and Toll pathway, BLASTx or BLASTp (version 2.2.31+) search were performed using the fruit fly, *Drosophila melanogaster* (Diptera; Meigen, 1830), sequences as query sequences from FlyBase (URL: https://flybase.org/ accessed October 1, 2022) and predicted amino acid sequence of *M. separata* as subject sequences. Phylogenetic tree construction (Neighbor join (NJ) method with jacard methods) and domain analysis of Peptidoglycan recognition proteins (PGRPs) and Gram-negative binding protein (GNBP) (hmmscan version 3.1b2 with Pfam-A.hmm of Pfam_35) were performed by DoMosaics (URL: https://domainworld.uni-muenster.de/developing/domosaics/ version 0.95) [44] while other phylogenetic trees of PGRPs and GNBPs (Maximum Likelihood method (ML)) were constructed by MEGA X with default settings [45]. Phylogenetic tree construction (Neighbor join method) and domain analysis of CTLswere performed by DoMosaics as same settings of the analyzing PGRPs and GNBPs. SignalP (version 6.0) and DeepTMHMM (version 1.0) were used to search for signal peptide sequences and transmembrane domains in CTLs, respectively [46,47].

## 3. Results

### 3.1. Construction of the genome sequences of M. separata

Using long-read and short-read genome sequence data of *M. separata*, we constructed *de novo* genome sequences. The genome construction scheme is shown in Figure 1. The assembled genome sequence data were first constructed using only the long-read sequence data, with a mean coverage of 187_X_. Subsequently, the assembled genome sequence data was polished using short-read sequence data. The basal status of the polished genome sequence data is shown in Table 1. The total length of the polished genome data is not unusual as a lepidopteran genome with a high contig N50 value (2,745,150 bp), considering the genome sizes of lepidoptera species. Subsequently, BUSCO assessed the completeness of the polished genome data (Table 2). Although most of the core genes in the BUSCO dataset were found in the polished genome data (Complete BUSCO), some core genes, which are single-copy genes, were found to be duplicated. The reason for the duplications could be the high regional heterogeneities in diploid genomes, which eventually led to incorrect assembly of two contigs that were assembled as one contig [48]. Therefore, to address this problem, the polished genome data were loaded into the Purge Haplotigs. The improved genome data from Purge Haplotigs (referred to as the finished genome data in Table 2) were analyzed using BUSCO (Table 2). Almost all core genes in the eukaryota_odb10 or lepidoptera_odb10 data set were found to be complete and single-copy BUSCO in the improved genome data, indicating that the genome data had good qualities as reference genome data, with a CG percentage of 38.60%. Hereafter, we refer to these genome data as the reference genome data or simply “the genome data” (deposited in DDBJ/ENA/GenBank; see Material and Methods 2.2).

**Table 1.**
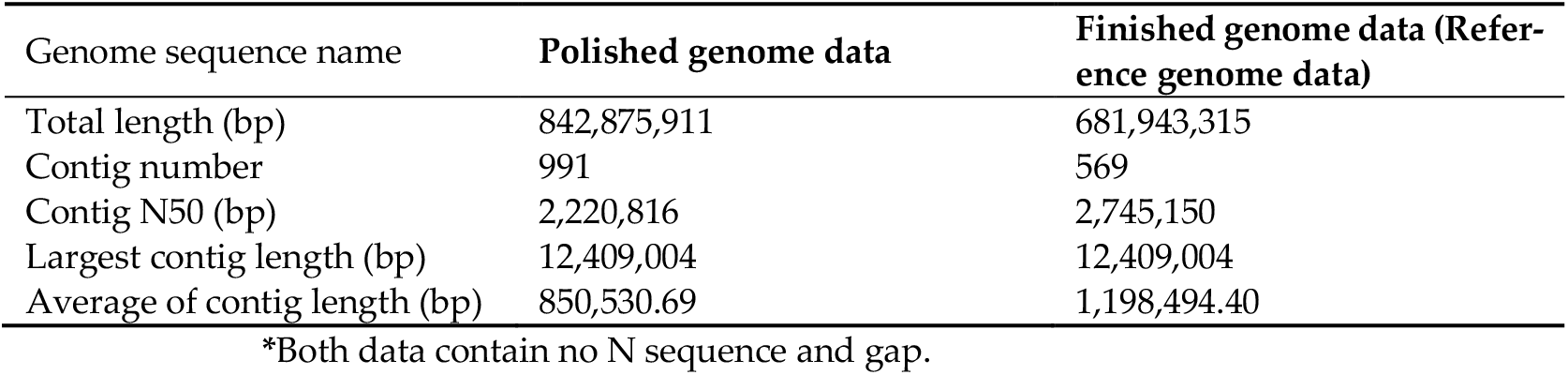
The status of the constructed genome sequence data*

| Genome sequence name | Polished genome data | Finished genome data (Reference genome data) |
| --- | --- | --- |
| Total length (bp) | 842,875,911 | 681,943,315 |
| Contig number | 991 | 569 |
| Contig N50 (bp) | 2,220,816 | 2,745,150 |
| Largest contig length (bp) | 12,409,004 | 12,409,004 |
| Average of contig length (bp) | 850,530.69 | 1,198,494.40 |

\*Both data contain no N sequence and gap.

**Table 2.**
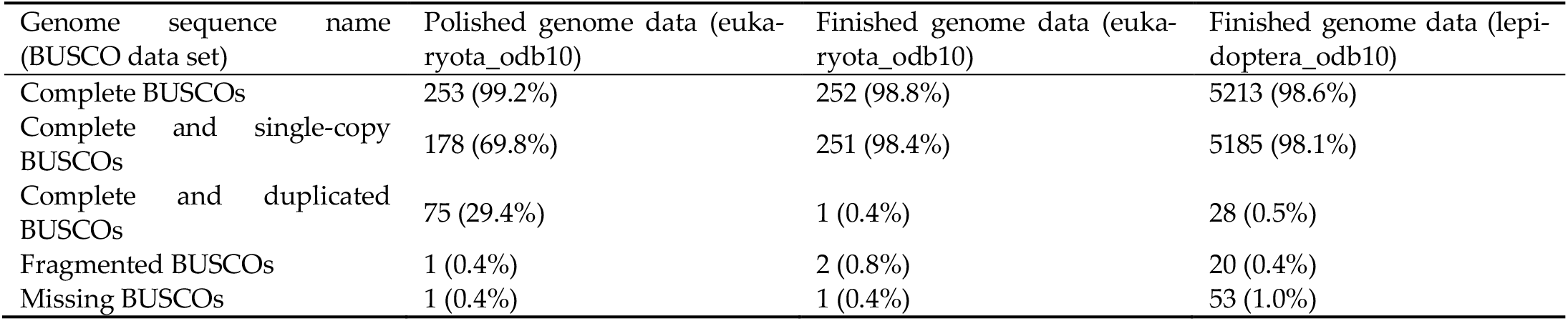
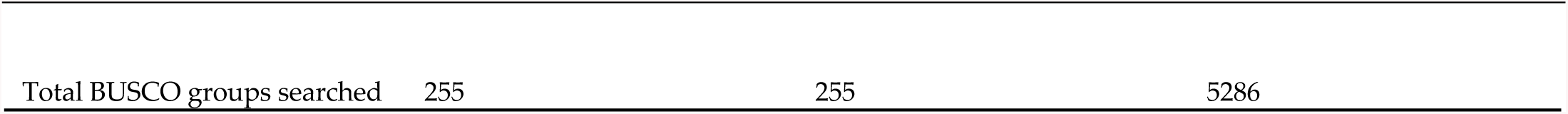
The results of BUSCO using the constructed genome sequence data

### 3.2. Search for Repetitive region and Gene prediction

We searched for the repetitive regions and transposable elements (TEs) in the genome sequence (Figure 1). First, *de novo* TE detection was performed, and 1,905 consensus sequences of TEs were constructed (Supplemental Data 3). TEs and repetitive regions in the *M. separata* genome were identified using consensus sequences; the summary status is shown in Table 3, and the output files are provided in Supplementary Data 4. Approximately 46.59% of the *M. separata* genome was repetitive or contained TE sites. Among the annotated TEs, the number of retroelements (Class I TEs) was much larger than that of DNA transposons (Class II TEs). Long interspersed nuclear elements (LINEs) were the most abundant retroelements, whereas Tc1-IS630-Pogo was the most abundant DNA transposon. Rolling circles covered 6.86% of the *M. separata* genome, whereas satellites and simple repeats were less than 1%.

**Table 3.** Transposons and repetitive regions in *M. separata* genome.

| total length: | 681943315 bp |  |  |
| --- | --- | --- | --- |
| bases masked: | 317706294 bp ( 46.59 %) |  |  |
|  | Number of elements* | length occupied | Percentage of sequence |
| Retroelements | 483,943 | 102,500,601 bp | 15.03% |
| SINEs: | 69 | 3,659 bp | 0.00% |
| Penelope | 0 | 0 bp | 0.00% |
| LINEs: | 466,443 | 94,162,645 bp | 13.81% |
| CRE/SLACS | 11,265 | 2,482,075 bp | 0.36% |
| L2/CR1/Rex | 47,274 | 15,582,975 bp | 2.29% |
| R1/LOA/Jockey | 125,283 | 28,136,850 bp | 4.13% |
| R2/R4/NeSL | 8,079 | 2,985,258 bp | 0.44% |
| RTE/Bov-B | 184,015 | 31,548,956 bp | 4.63% |
| L1/CIN4 | 0 | 0 bp | 0.00% |
| LTR elements: | 17,431 | 8,334,297 bp | 1.22% |
| BEL/Pao | 1,963 | 3,457,738 bp | 0.51% |
| Ty1/Copia | 2,715 | 1,358,093 bp | 0.20% |
| Gypsy/DIRS1 | 6,472 | 2,769,678 bp | 0.41% |
| Retroviral | 160 | 76,839 bp | 0.01% |
| DNA transposons | 80,267 | 20,701,313 bp | 3.04% |
| hobo-Activator | 6,121 | 1,085,735 bp | 0.16% |
| Tc1-IS630-Pogo | 24,800 | 10,838,060 bp | 1.59% |
| En-Spm | 0 | 0 bp | 0.00% |
| MuDR-IS905 | 0 | 0 bp | 0.00% |
| PiggyBac | 166 | 236,668 bp | 0.03% |
| Tourist/Harbinger | 2,055 | 550,091 bp | 0.08% |
| Other (Mirage, P-element, Transib) | 110 | 177,033 bp | 0.03% |
| Rolling-circles | 281,101 | 46,800,852 bp | 6.86% |
| Unclassified: | 880,600 | 14,0919,871 bp | 20.66% |
| Total interspersed repeats: |  | 26,4121,785 bp | 38.73% |
| Small RNA: | 204 | 23,041 bp | 0.00% |
| Satellites: | 177 | 69,651 bp | 0.01% |
| Simple repeats: | 102,349 | 5,964,291 bp | 0.87% |
| Low complexity: | 15,392 | 726,674 bp | 0.11% |

\* most repeats fragmented by insertions or deletions have been counted as one element

**Table 4.** Numbers of the hit genes in each data set by Fanflow4

|  | Numbers of the hit proteins (among 24,453 proteins) | Percentage of the hit proteins (%) |
| --- | --- | --- |
| *H. sapience* | 14,037 | 57.4039995 |
| *M. musculus* | 13,699 | 56.021756 |
| *C. elegans* | 11,235 | 45.9452828 |
| *D. melanogaster* | 12,879 | 52.6683842 |
| *B. mori* | 19,704 | 80.5790701 |
| *M. sexta* | 17,996 | 73.594242 |
| *A. mellifera* | 14,427 | 58.9988958 |
| *T. castaneum* | 16,496 | 67.4600254 |
| Unigene | 16,086 | 65.7833395 |
| Pfam | 13,244 | 54.1610436 |

Gene predictions of *M. separata* were performed using the masked genome data (Supplementary Data 1) and RNA-Seq data as hint data, of which some were retrieved from SRA public database (Supplementary Data 1 and Figure 1). A total of 21,970 genes were identified in the *M. separata* genome (gene.gtf in Supplementary Data 5), whereas 24,452 coding sequences (CDSs) and amino acid sequences were predicted because some genes had multiple transcripts (gene_cds.fasta and gene_pep.fasta in Supplementary Data 5). Using the predicted amino acid sequences, the predicted genes were functionally annotated with Fanflow4Insects (Supplementary Data 6)[43]. Approximately 45–57% of the protein sequences were annotated using gene sets of model species (*Homo sapiens, Mus musculus*, and *Caenorhabditis elegans* (Maupas, 1900)). Among the insect species datasets, a relatively higher percentage of protein sequences were annotated using Lepidoptera datasets (*M. sexta* and *B. mori*) than using other insect species datasets (*D. melanogaster*, the western honeybee (*Apis mellifera*; Hymenoptera; Linnaeus, 1758), and the red flour beetle (*Tribolium castaneum*; Coleoptera; Herbst, 1797)), and over half of the sequences were annotated using Pfam and Unigene datasets.

We checked whether the microbial sequences were contaminated in the reference genome and attempted to remove the microbial sequences using annotation data of the gene set with NCBI-nr (Supplementary Data 7). We counted the numbers of the names of the microbial species in the annotation data (Supplementary Data 7). A total of 32, 23, 39, 12, and 10 genes were annotated with orthologs from *Pseudomonas aeruginosa, Piscirickettsia salmonis*, Spodoptera moth adenovirus 1, *Trichoplusia ni* TED virus, and *Conidiobolus coronatus*, respectively. If these genes are located in the same contig and the contig sequences have sequences identical to those in the microbial species, the corresponding contigs needed to be removed from the reference genome. The gene sets annotated with *P. aeruginosa, P. salmonis*, Spodoptera moth adenovirus 1, and Trichoplusia ni TED virus descriptions were not located in the same contigs. Ten genes annotated with the *C. coronatus* descriptions were located in the contig; however, this contig did not hit any genome sequences of *C. coronatus*. Therefore, we did not remove any contigs from the *M. separata* reference genome sequences.

### 3.3. Immune-related genes in M. separata

To determine whether antimicrobial peptide (AMP) production systems function in *M. separata*, we searched for genes in the Toll and IMD pathways, which have been thoroughly investigated in *D. melanogaster* [19]. Using data sets of the Toll and IMD pathway sequences in FlyBase, we determined whether orthologs of the core components of the Toll and IMD pathways exist in *M. separata*. Orthologs of almost all core intracellular components of the Toll and IMD pathways were found in *M. separata* (Figure 2). Although *IMD* and *MyD88* were not found by BLASTx (Supplementary Data 8), both genes were identified by BLASTp (Supplementary Data 9).

**Figure 2.**
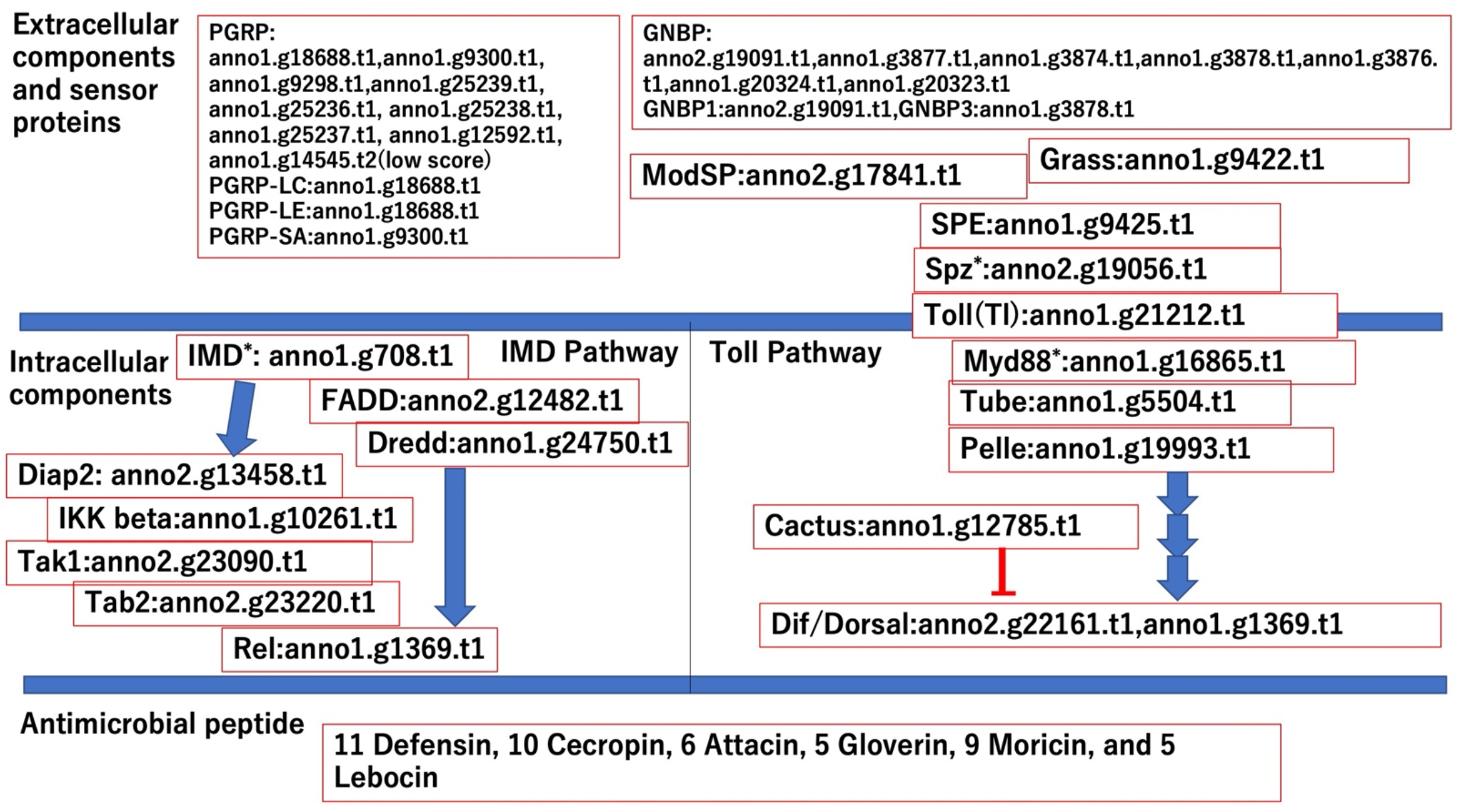
Gene IDs of the orthologues consisting of Toll and IMD pathway based on BLASTx search. Number of genes of antimicrobial peptides were shown in bottom of this figure, and the gene IDs of the Orthologs of antimicrobial peptide are shown in Supplemental Data 10. Asterisks indicates that the orthologues were found by BLASTp search.

Extracellular components in Toll pathways were searched [19]. Orthologues of *ModSP, Grass* and *Spaezle-processing enzyme* (*SPE*), which are forming serine protease cascade between sensor proteins and Toll receptor complex, were found in *M. separata* genes while *Toll* (*Tl*) and *Spaezle* (*Spz*) (found by the BLASTp search) orthologs consisting of the Toll receptor complex were found (Figure 2). Sensor proteins of IMD and Toll pathway recognize the surface structures of the infected microbes and arouse signals to activate these pathways. PGRPs recognize the peptidoglycan structure derived from gram-positive or gram-negative bacteria. BLASTx results showed that nine genes in the *M. separata* genome were orthologs of PGRPs; orthologs of *PGRP-LE* and -*LC* (the IMD pathway) and *PGRP-SA* were found, and the genes showing minimum e-values among *M. separata* nine PGRPs in the BLASTx results were allocated as orthologue of each three PGRP (Figure 2). GNBPs also acts as a sensor protein for fungi and gram-positive bacteria. Seven GNBP-encoding genes were found in the *M. separata* genome, whereas *D. melanogaster* harbored only three such genes. Orthologs of GNBP1 and GNBP3 showing minimum e-values among *M. separata* GNBPs in the BLASTx results were allocated (Figure 2). Furthermore, the NJ phylogenetic trees of PGRPs and GNBPs were constructed, and the domain analysis was performed by hmmerscan bundled in DoMosaics, and the ML phylogenetic trees of PGRP and GNBP were constructed (Supplemental Data 10). In the PGRP NJ tree, three *D. melanogaster* PGRPs and *M. separata* anno1.g18688.t1 (annotated PGRP-LC and LE by BLASTx results.) formed a clade while *M. separata* anno1.g9300.t1 belonged to another clade. The domain analysis of the PGRP showed that most of *M. separata* PGRPs possessed Amidase_2 domain, which *D. melanogaster* PGRPs harbored. The structure of the ML tree of PGRP were different from that of the NJ tree of PGRP. In the ML tree, *D. melanogaster* PGRP-LC and *D. melanogaster* PGRP-SA formed a clade with anno1.g9298.t and anno1.g14545.t, respectively while *D. melanogaster* PGRP-LE did not form a clade with a single *M. separata* PGRP. In the GNBP NJ tree, anno1.g3877.t1 and anno1.g3876 formed the same clades of *D. melanogaster* GNBP1 and GNBP3, respectively, and both two *M. separata* GNBP possess CBM39 and Glyco_hydro_16 domain, which *D. melanogaster* GNBPs harbored. The structures of the ML tree were different from that of the NJ tree. *D. melanogaster* GNBP1 formed a clade with two *M. separata* GNBPs (anno1.g20324 and anno1.g20324), which are considered as paralogous genes, while *D. melanogaster* GNBP3 did not form a clade with a single *M. separata* GNBP. Taken together, there are different relationships of orthologues between *D. melanogaster* and *M. separata* in the NJ and ML trees. Furthermore, there are inconsistencies between the BLASTx and phylogenetic tree results. The reasons of them might be because the methods of calculations of the sequence similarities were different, and the degrees of differences within PGRPs and GNBPs sequences might not be so high, especially, in PGRPs (low bootstrap values in the ML tree). The functional analysis of these proteins of *M. separata* will be required in future research.

AMPs are effector molecules against invading microbes, and the induction of AMP genes is regulated mainly by the Toll and IMD pathways in *D. melanogaster* [49]. Defensin, cecropin, attacin, gloverin, moricin, and lebocin are the major AMP genes in insects [50]. Using *M. separata* gene dataset and annotation results (Supplementary Data 6), orthologs of the AMP genes in *M. separata* were searched (Supplementary Data 11). This is because most AMPs are very short, and orthologs cannot be found through BLAST alone. Although orthologs of defensin and cecropin were not found with low E-values either by BLASTp or hmmer domain search (data not shown), 11 defensins and 10 cecropins were found in the annotation data of *M. separata*. The 11 defensins were annotated with “Defensin*”* descriptions of several insect species such as *D. melanogaster* and *Aedes aegypti* (“UniGene-description” column in Supplemental Data 6), whereas the 10 cecropins were annotated with “Cecropin” descriptions of *M. sexta*. Six attacins, five gloverins (anno2.g16145.t1 was also annotated as attacin), nine moricins (seven moricins were also annotated as cecropins), and five lebocins were found in *M. separata*.

As described above, CTLs recognize a wide range of ligands [51,52]. Therefore, CTLs are involved in several immune reactions of insects by binding to the surfaces of not only the invaded microbes and organisms but also their own cells and tissues. CTLs possess more than one C-type lectin domain (CTLD) (also known as the “carbohydrate recognition domains”; CRD). Thus, CTL genes in *M. separata*, which were annotated as possessing CTLD (Pfam ID: PF00059), were searched. Among *M. separata* genes, 105 CTLs were identified (Supplemental Data 12). The results of the domain analysis by hmmerscan bundled in DoMosaics and sequences analysis of the 105 CTL are shown in Supplemental Data 13. Most CTL possessed two CTLDs classified as Dual CTLD, and 12 and 6 possessed single and more than three CTLDs, respectively (Table 5). Five CTLs possessed CTLD and other domains, which were classified as CTL-X. Furthermore, we determined whether the CTLs possessed signal peptide sequences and transmembrane domains. Of the 105 CTLs, 77 and 84 were predicted to possess signal peptides, using SignalP and DeepTMHMM, respectively (Table 5). This difference in the number of predicted signal peptides may be because of the difference in the prediction algorithms of the two software. These four CTLs possessed a transmembrane domain.

**Table 5.**
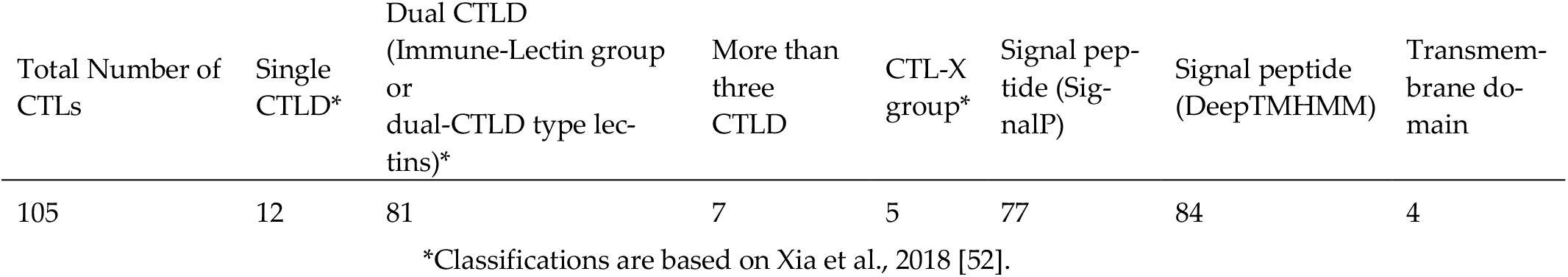
Summary of CTL analyses

## 4. Discussion

In this study, the reference genome sequences and gene dataset of the oriental armyworm, *M. separata*, were constructed. Using long-read and short-read genome data, genome sequences with high contig N50 values and good BUSCO scores were constructed through polishing and purging haplotig processes. After searching for repetitive regions and TEs in the genome sequences and masking such sequences, we obtained gene set data, using masked genome data and RNA-Seq data, some of which were retrieved from SRA. Consequently, the gene set was functionally annotated, and using amino acid sequence data of the gene set and the functional annotation data, orthologs of the IMD and Toll pathways as well as CTLs, which are involved in immune reactions in model insect species, were identified.

The contig N50 value of the reference genome we constructed was approximated at 2.7Mbp, and the results of the BUSCO analysis showed that over 98 % of BUSCO genes (core genes) were identified. These results suggest that the genome sequences data of *M. separata* possess sufficient quality as the reference genome in term of continuity and completeness of genome, which can be used for the various studies, especially genome or molecular research. The size of *M. separata* finished genome sequence was approximately 628 Mbp, as against these of *P. xylostella, T. ni, B. mori, S. litura, S. exigua*, and *M. sexta* that were 328 Mbp, 333Mbp, 460 Mbp, 438 Mbp, 419 Mbp, and 470 Mbp, respectively [2,9,12,13,15,17]. The genome size of butterflies (264 species) varied from 195 to 1262 Mbp [18]. Considering these results, the genome size of *M. separata* is larger than those of the other many lepidopteran species except those of butterflies, and but it is a reasonable in the range of some lepidopteran insects.

Further, it was found that approximately 48% of the *M. separata* genome contained repetitive regions or TEs, and retroelements were the major TEs, of which LINEs were the major components. In *B. mori*, approximately 46.5% of the genome contained repetitive regions or TEs, and more Class I retroelements than Class II DNA elements were detected, which is the same tendency as in the *M. separata* genome [9]. Additionally, SINEs consist of approximately 12% of the *B. mori* genome, and LINEs consist of approximately 17%, which are comparable numbers. However, approximately 32% of the *S. litura* genome and more Class I retroelements (about 11%) than Class II DNA elements (about 2%) were detected. LINEs and SINEs consist of about 8% and 2% of the *S. litura* genome, respectively [2]. In *A. mellifera*, a hymenopteran model species, the total repetitive region consists of approximately 11% of the genome, and Class II DNA elements are major components of the TEs, which are clearly different from lepidopteran families [53]. Taken together, these results suggest that the comprehensive tendency of *M. separata* was the same as the model lepidopteran species, and detailed status, especially the ratio of SINEs were different, which may be *M. separata-*specific features of TEs.

A total of 21,970 genes (24,452 CDSs) were predicted in the *M. separata* genome, whereas *B. mori, S. litura, S. exigua*, and *M. sexta* had 16,880 genes, 15,317 protein-coding genes, 18,477 transcripts, and 25,256 genes, respectively [2,9,12,17]. The differences among the species may be partially owing to the methods used to prepare the gene sets. Considering these results, the number of genes of *M. separata* may be reasonable for lepidopteran gene sets. The annotation ratios of *H. sapiens, M. musculus*, and *C. elegans* were lower than those of *T. castaneum, M. sexta*, and *B. mori*. The ratio of *D. melanogaster* to *A. mellifera*, a model insect species, was not very high. According to a phylogenetic tree constructed by Misof et al. [54], Hymenoptera branched at the earliest time, Coleoptera branched, and Lepidoptera and Diptera branched. Our ratios were not consistent with this tree structure. Although we could not determine the reasons underlying these differences, the gene sets could provide new evolutionary insights in such research fields.

Searching for the genes consisting of the Toll and IMD pathways revealed that intracellular components of these pathways are conserved, which has been observed in multiple insect species [19,55–57], suggesting that AMPs are regulated by the two signaling pathways in *M. separata*. The numbers of PGRPs and GNBPs predicted to function as pattern recognition receptors (PRRs) for fungi and bacteria [58] were different from those in other insect species [19,55,57,59,60]. Nine PGRPs and seven GNBPs were identified in the *M. separata* genome. Among these genes, the orthologs of PGRP-LC, -LE and -SA, GNBP1, and GNBP3 function as PRRs in *D. melanogaster*. However, functional analysis using some non-*Drosophila* species showed that the orthologs did not play a pivotal role in sensing microbes [60,61], which led to different signal transduction in the Toll and IMD pathways in *D. melanogaster*. The results suggest that *M. separata* has the Toll and IMD pathway systems, which have different signal transduction flows from *D. melanogaster*, as observed in *Plautia stali* and *T. castaneum* [60,62]. Over 40 AMP genes were identified in the *M. separata* genome and included typical AMP genes, such as cecropin, defensin, and attacin. However, some of the AMP genes were annotated without typical domain annotations in the Pfam database analysis. For example, the cecropin domain was not identified in the 10 cecropin genes annotated by the *M. sexta* gene set. Thus, further investigation is warranted to determine whether such AMP genes actually exist or are expressed, using RNA-Seq or quantitative reverse transcription-PCR analysis.

According to Xia et al., the number of CTL genes in insects ranges from 4 to 40 [52]. Among lepidoptera species, *B. mori* and *M. sexta* have 23 and 34 CTL genes, respectively, whereas *P. xylostella* has only seven genes. Compared to these species, *M. separata* has an outstandingly higher number of CTL genes. Notably, approximately 90% of CTLs have two CTLDs (or CRDs), which were categorized as dual CTLD type CTLs and more CTLD-type CTLs. Additionally, approximately > 70% CTLs were found to have a signal peptide sequence, five CTLs were classified as CTL-X, and four CTLs had transmembrane domains, and these results are not different from those obtained for other insects [52]. Based on these results, we assumed that the dual-CTLD-type lectin genes were duplicated in the *M. separata* genome. Many dual-CTLD-type lectins regulate the immune system of insects. Immunlectin II, a dual-CTLD-type lectin from *M. sexta*, binds to bacterial lipopolysaccharide with a CTLD [63]. In *B. mori*, dual-CTLD-type CTLs are required for nodule formation [64]. EPL, which enhances encapsulation, is also a dual-CTLD-type lectin [21]. Therefore, dual-CTLD-type lectins are expected to play more important roles in immunity in *M. separata* than in other insects. Recently, Sawa et al. reported that the parasitoid wasp *Cotesia cariyai* suppressed melanization and encapsulation of *M. separata* by manipulating the CTLs of both species, suggesting that CTLs are key factors for the success of parasitization [65]. To shed light on the well-controlled immune system and its interaction with various microorganisms and parasitoids, analyzing the diversified lectins and the molecules they recognize will be necessary.

## 5. Conclusions

We constructed the reference genome of *M. separata* (size: 682 Mbp) with sufficient qualities as the reference genome, in terms of continuity and completeness of core genes. In addition, 21,970 genes were predicted using genome sequence data, and functional annotations were performed. Using the gene set and annotation data, several immune-related genes were identified, providing new insights into the genomics and immunology of *M. separata*. We consider that the obtained reference genome data can promote studies on molecular biology of *M. separata* and comparative genomics of insects in general.

## Supplementary Materials

All Supplemental materials are available in figshare. For the detail, See “Data Availability Statement” section.

## Author Contributions

Conceptualization, K.Y., S.F. and H.B.; methodology, K.Y., A.J. and H.B.; validation, K.Y. A.J. and H.B.; formal analysis, K.Y. and H.B.; resources, S.F. and R.Z; data curation, K.Y., S.F., A.J. and H.B; writing—original draft preparation, K.Y.; writing—review and editing, K.Y., S.F., A.J. and H.B; visualization, K.Y.; supervision, K.Y.; project administration, K.Y.; funding acquisition, K.Y. and S.F. All authors have read and agreed to the published version of the manuscript.

## Institutional Review Board Statement

Not applicable.

## Informed Consent Statement

Not applicable.

## Data Availability Statement

All Supplemental data are available in figshare. DOI: 10.6084/m9.figshare.c.6192784.

**Supplemental Data 1** Sequence Read Archive accession IDs of raw sequence data in this study. DOI: 10.6084/m9.figshare.21082570.

**Supplemental Data 2** DDBJ/ENA/GenBank accession IDs and FASTA file IDs of assembled genome sequences. DOI: 10.6084/m9.figshare.21215255.

**Supplemental Data 3** Consensus sequences of transposable elements in *M. separata*. The consensus sequences of transposable elements in *M. separata* were constructed by RepeatModeler2. The sequence files are FASTA file and stk file from RepeatModeler2. DOI: 10.6084/m9.figshare.21230882

**Supplemental Data 4** Output files of RepeatMasker. Armyworm_genome_seq_finished.fa.out shows the detailed status of all detected transposable elements (TEs) and their repetitive positions. Armyworm_genome_seq_finished.fa.cat.gz shows the sequence comparison between all TEs and genome sequences. Armyworm_genome_seq_finished.fa.masked.gz is the genome sequence in which repetitive and TE sequences are indicated in lowercase letters. See RepeatMasker for detailed explanation (https://www.repeatmasker.org/). DOI: 10.6084/m9.figshare.21231500.

**Supplemental Data 5** Data related to predicted genes, coding sequences and amino acid sequences in *M. separata* by Braker2. Gene.gtf contains structural data of predicted genes and transcripts. Gene_cds.fasta and gene_pep.fasta contain predicted coding sequences and amino acid sequences respectively. DOI: 10.6084/m9.figshare.21257601.

**Supplemental Data 6** Functional annotations of *M. separata* genes. Using the predicted amino acid sequences of *M. separata*, its genes were functionally annotated using Fanflow4Insects. The predicted amino acid sequences were compared with the gene set data of *Homo sapiens, Mus musculus, Caenorhabditis elegans, Drosophila melanogaster, Bombyx mori, Manduca sexta, Apis mellifera, Tribolium castaneum*, and UniProtKB/Swiss-Prot dataset. “pid” and “gene_symbol” indicate hit protein IDs and gene symbols of each dataset, respectively. The protein domains were annotated using Pfam and HMMER. DOI: 10.6084/m9.figshare.21257625.

**Supplemental Data 7** BLASTp annotation results of the predicted gene set against the NCBI-nr database and count results of the number of species that annotate *M. separata* genes. Input _pep.fasta.summary is the result of annotation of *M. separata* genes against the BLAST_nr database. results.txt contains counting results for the number of species that annotate *M. separata* genes. DOI: 10.6084/m9.figshare.21386523.

**Supplemental Data 8** BLASTx results for genes in the Toll and IMD pathways. BLASTx was performed using either *D. melanogaster* transcript sequences consisting of the Toll (ID: FBgg0001194) or IMD (ID: FBgg0001059) pathway components in Flybase as query sequences and *M. separata*’s predicted amino acid sequences as subject sequences. DOI: 10.6084/m9.figshare.21325707.

**Supplemental Data 9** BLASTp results for genes in the Toll and IMD pathways. BLASTp was performed using either *D. melanogaster* amino acid sequences consisting of the Toll (ID: FBgg0001194) or IMD (ID: FBgg0001059) pathway components in Flybase as query sequences and *M. separata*’s predicted amino acid sequences as subject sequences. DOI: 10.6084/m9.figshare.21325740.

**Supplemental Data 10** Phylogenetic trees and domain analysis of PGRPs and GNBPs of *M. separata* and *D. melanogaster* using DoMosaics and MEGA X. NJ phylogenetic trees of GNBPs and PGRPs, plus the domain structure of each gene by DoMosaics, and ML phylogenetic trees by MEGA X with bootstrap values are shown in PGRP_GNBP_tree.pptx. PGRP.nwk and GNBP.nwk are the files used for the construction of a phylogenetic tree of DoMosaics. PGRP.domtree and GNBP.domtree are the raw files of the DoMosaics phylogenetic trees. Two mtsx files were session files of ML trees of PGRP and GNBP for MEGA X. Genbank accession IDs of *D. melanogaster* PGRP and GNBP are list in Accession_IDs_Dm_GNBP_PGRP.csv. DOI: 10.6084/m9.figshare.21709790.

**Supplemental Data 11** ID list of orthologs of the six antimicrobial peptide (AMP) genes. Orthologs of the six AMP genes (defensin, cecropin, attacin, gloverin, moricin, and lebocin) in *M. separata* were searched using the annotation results (Supplementary Data 6). The *M. separata* gene set IDs of the AMP orthologs are listed. DOI: 10.6084/m9.figshare.21346908.

**Supplemental Data 12** Domain and phylogenetic analysis of 105 C-type lectins identified in *M. separata*, using DoMosaics. CTL_tree_domain.jpg is the CTL tree developed using DoMosaics. C_type_lectin_ID.txt shows the ID of C-type lectin genes in *M. separata*, and CTL.nwk is a new file for the construction of a phylogenetic tree. The CTL.domtree file is the raw file of the CTL_tree_domain.jpg. DOI: 10.6084/m9.figshare.21351501.

**Supplemental Data 13** SignalP results for signal peptide sequences and DeepTMHMM results for the transmembrane domain. output_signalP.gff3 and SignalP_results.txt are the output files of SignalP, and DTU_DeepTMHMM_1.0.15-results.zip is the output file of DeepTMHMM. For details, please refer to the references of the software. DOI: 10.6084/m9.figshare.21436560.

## Acknowledgments

We deeply acknowledged Prof. Dr. Atsushi Toyoda at Advanced Genomics Center, The National Institute of Genetics, Mishima, Shizuoka, Japan for the sequencing the prepared gDNA samples.

